# The experimentally evolved fluconazole-resistant Clade II isolates of *Candidozyma auris* exhibit a distinct lipid compositional landscape, highlighting intra-clade sphingolipid heterogeneity

**DOI:** 10.1101/2025.03.13.642928

**Authors:** Praveen Kumar, Basharat Ali, Mohit Kumar, Hans Carolus, Celia Lobo Romero, Rudy Vergauwen, Anshu Chauhan, Aswathy Narayanan, Atanu Banerjee, Naseem A. Gaur, Ashutosh Singh, Patrick Van Dijck, Arunaloke Chakrabarti, Shiva Prakash M. Rudramurthy, Kaustuv Sanyal, Rajendra Prasad

## Abstract

The intrinsic resistance of *Candidozyma auris* to antifungal drugs poses a major therapeutic challenge, with conventional resistance mechanisms providing only partial explanations. Sphingolipids (SLs), known for their interclade heterogeneity, play a crucial role in antifungal resistance. This study examined the SL landscape in two drug-susceptible clade II isolates, C-line and P-line, from distinct geographical origins, which were experimentally evolved to develop stable fluconazole (FLC) resistance. The progenitors displayed distinct SL profiles, P1 had higher PhytoCer and αOHPhytoCer, indicating a more active acidic SL biosynthesis branch, whereas C1 exhibited elevated αOHGlcCer, αOHCer, and LCBs, reflecting a greater role of the neutral biosynthesis branch. The principal component analysis (PCA) also confirmed distinct segregation of the two progenitors. Upon evolution, P1.1 and C1.1 adaptors showed significant SL alterations. P1.1 exhibited PhytoCer enrichment, while C1.1 showed reduced αOHGlcCer alongside increased PhytoCer, dhCer and αOHPhytoCer levels. Notably, αOHGlcCer remained unchanged in P1.1, whereas LCBs and αOHPhytoCer decreased compared to P1. Despite these lineage-specific differences between the progenitors, both evolved replicates exhibited increased PhytoCer as a common denominator like what is also observed in clinical FLC-resistant isolates. These findings highlight intra-clade SL variability and suggest that specific SLs contribute to FLC resistance in *C. auris*.

## Introduction

The emergence of multidrug-resistant *Candidozyma auris* across five continents has raised significant global concern. These primarily clonal clades spread rapidly within healthcare settings and persist on surfaces and bedding (Berkow *et al*. 2017; Kordalewska and Perlin 2019). A multicentred study spanning three continents reported that approximately 93% of *C. auris* isolates were resistant to FLC, around 35% to amphotericin B (AmB), 41% resistant to both FLC and AmB, and about 4% resistant to all three major antifungal classes, including FLC, AmB, and echinocandins (Lockhart and KA 2016a). Azoles, frontline drugs for systemic infections caused by most *Candida* species, target the ergosterol biosynthesis pathway by inhibiting 14α-demethylase, encoded by *ERG11* (Lupetti *et al*. 2002). Early analyses of drug-resistant *C. auris* isolates identified mutations in the azole target *ERG11* as key contributors to resistance, findings supported by whole-genome sequencing (Lockhart and KA 2016b; Muñoz *et al*. 2018; Kim *et al*. 2019; Rybak *et al*. 2019). Additionally, phenotypic studies revealed the potential overexpression of drug transporter genes, such as *CDR1*, *CDR2*, and *MDR1*, which enhances drug efflux (Chatterjee *et al*. 2015; Kim *et al*. 2019; Rybak *et al*. 2019). Notably, there are instances where none of the established mechanisms, such as *ERG11* mutations or drug efflux, appear to play a dominant role in resistance (Rybak *et al*. 2020; Narayanan *et al*. 2022). Collectively, these findings suggest that well recognized resistance mechanisms alone cannot fully explain the high intrinsic resistance observed in *C. auris*.

While conventional antifungal drugs such as azoles and polyenes primarily target ergosterol, alterations in SL classes suggest that these biomolecules, along with ergosterol, are pivotal lipid components that interact within the membrane domain (Song, Liu and Li 2020). This connection is further supported by earlier observations in *C. albicans*, where the deletion of specific SL biosynthesis genes increased susceptibility to antifungals and disrupted the localization of major multidrug exporter proteins, Cdr1p (Pasrija, Panwar and Prasad 2008; Gao 2018). Several studies have established that SLs contribute to azole resistance (Gao 2018; Garnaud *et al*. 2018) and function as signalling molecules with critical roles in various cellular processes (Chen *et al*. 2013; Montefusco *et al*. 2013; Huang, Withers and Dickson 2014; Rego *et al*. 2014; Teixeira and Costa 2016). Similarly, long-chain bases (LCBs), key intermediates in SL biosynthesis including dihydrosphingosine (DHS), phytosphingosine (PHS), and their phosphorylated forms (DHS-1-P and PHS-1-P) are involved in intracellular signalling in *Saccharomyces cerevisiae* and *C. albicans* (Kobayashi and Nagiec 2003; Vandenbosch *et al*. 2012). Additionally, specific SL molecules, such as glucosylceramide (GlcCer) and inositol phosphorylceramide (IPC), contribute to fungal pathogenesis and virulence (Nimrichter and Rodrigues 2011).

Despite extensive evidence supporting the roles of SLs and membrane sterols in azole resistance, similar studies in *C. auris* remain limited. Recently, a comparative lipidomic analysis between drug-resistant and susceptible *C. auris* isolates revealed a significant remodelling of polar lipids in drug-resistant strains (Shahi *et al*. 2022). Similarly, Kumar et al. (2021) characterized the molecular SL signatures of drug-resistant clinical *C. auris* isolates from Indian hospitals and demonstrated their susceptibility to SL pathway inhibitors, underscoring the importance of SLs in drug susceptibility. More recently, Ali *et al* reported interclade heterogeneity in SL composition among *C. auris* isolates from different geographical regions (Ali *et al*. 2024b). While these studies highlight the relevance of SLs in drug resistance, they do not clarify how SL modulation influences the development of resistance. Recent experimental evolution studies, including ours, have mapped adaptive trajectories that reveal the emergence of antifungal resistance through multiple pathways, ultimately leading to FLC resistance in *C. auris* (Narayanan *et al*. 2022). The experimentally evolved FLC-resistant adaptors provided a unique opportunity to investigate the role of SLs in modulating azole resistance. In this study, two independently evolved azole-resistant adaptive lines of *C. auris* from clade II underwent high-throughput lipid analysis, which revealed distinct SL and ergosterol profiles in the evolved FLC-resistant lines. These findings underscore the intra-clade heterogeneity between FLC-susceptible and resistant *C. auris* strains.

## Materials and Methods

### Strains, culture media and growth conditions

*C. auris* clade 2 isolate CBS10913T was obtained from the Central Bureau voor Schimmel Cultures (CBS), The Fungal Biodiversity Centre of the Royal Netherlands Academy of Arts and Sciences (KNAW), Utrecht and NCCPF 470296 from National Culture Collection of Pathogenic Fungi (NCCPF) Post-Graduate Institute of Medical Education and Research (PGIMER), Chandigarh. FLC evolved isolates were the derivatives of the evolution experiment and stored at −80°C in 30% glycerol and grown in yeast extract peptone dextrose (YPD) at 30°C when revived. All media components and chemicals used were molecular grade and sourced from commercial suppliers.

### Experimental evolution of C. auris

The strains of *C. auris* CBS10913T (C-line) (MIC_50_ - FLC-8 μg/mL; AmB-0.5 μg/mL; and caspofungin (CSF-0.1 μg/mL) and NCCPF 470296 (P-line) (MIC_50_, FLC-16 μg/mL; AmB-0.5 μg/mL; and caspofungin (CSF-0.1 μg/mL) were employed for the experimental evolution. The experimental evolution protocol earlier reported (Gerstein and Berman 2020) was followed with slight modifications (Narayanan *et al*. 2022). Briefly, on the YPD plate, the strain was revived from a frozen stock, and cultured for 48 hours at 30°C. After that, a single colony was patched onto a fresh YPD plate and incubated at 30°C for 48 hrs. This single colony was inoculated in 10 mL of fresh YPD broth and cultured at 30°C, 180 rpm for 72 hours. Three separate tubes, each holding 9,990 μL of fresh YPD with or without FLC for both C and P-line (8 μg/mL and 16 μg/mL respectively), were filled with 10 μl of cells from the saturated culture with an optical density (OD_600_) of 0.1. The cultures were then cultured for an additional 72 hours at 30°C for 10 generations. A total of 10 such transfers were done in the presence and absence of FLC to obtain an evolved population after approximately 100 generations. Every 10 generations, cell aliquots were harvested and stored at −80°C in 30% glycerol for further experiments (Narayanan *et al*. 2022).

### Determination of minimum inhibitory concentration (MIC)

*C. auris* progenitor and adaptor lines were cultured in YPD at 30°C for overnight, and MIC tests were carried out in accordance with Clinical and Laboratory Standards Institute (CLSI) recommendations (*M27 Ed4 Broth Dilution Antifungal Susceptibility, Yeasts*). For an OD_600_ of 0.1, the cells were diluted in 0.9% saline solution. After that, the cells were diluted 100 times in YPD media. Equal amounts of media with 2-fold drug concentrations were prepared in round-bottomed, 96-well microtiter plates followed by addition of diluted cell suspension. The plates were incubated at 30°C for 48 hrs and then OD_600_ nm was determined in each well with microplate reader (iMark Bio-Rad). OD_600_ nm reading was used to evaluate the MIC test endpoint, which was defined as the lowest fluconazole drug concentration that produced 50% inhibition of growth (MIC_50_) in comparison to the growth of drug free control.

#### Sphingolipids extraction

The strains were first cultured in liquid YPD media for overnight in an incubator shaker at 30°C. After that, a secondary culture with a starting OD_600_ of 0.1 in 50 ml was grown until it achieved an OD_600_ of 1, indicating the mid-log cell growth phase. Cultures of each strain were centrifuged at 4000 rpm for five minutes to get 5× 10^8^ cells and cell pellets were washed with sterile water twice. C17 Sphingosine (d17:1) and C17 Ceramide (d18:1, 17:0)(Avantipolar Lipids, AL, USA) were added as internal standards prior to the lysis. This was followed by cell lysis using glass beads (50 mg, 0.4-0.6 mm) in a Fastprep® system (MP Biomedicals, CA, USA). Lipids were extracted from the lysate using methods described by (Kumar *et al*. 2021; Ali *et al*. 2024b). The extracted SLs were dried in the presence of N2 and kept at −20°C until further analysis.

### Liquid Chromatography-Mass Spectrometry (LC-MS)

The isolated lipids were reconstituted in an organic buffer consisting of 0.2% formic acid and 1 mM ammonium formate in methanol. The mobile phase system used was a two-buffer setup with an aqueous buffer containing 0.2% formic acid and 2 mM ammonium formate. A sample volume of 5µL was injected by Auto-sampler and resolved on the C8 column (Waters, MA, USA). A combined flow rate of 300 μL per minute was maintained throughout in a gradient manner as described in our previous study(Ali *et al*. 2024a). SLs were identified using the targeted multiple reaction monitoring (MRM) method in the QTRAP® 4500 mass spectrometer (SCIEX USA) as previously described(Kumar *et al*. 2021)

### Protein estimation

Protein estimation of the cell lysate was done using Bicinchoninic Acid (BCA) Protein Assay kit (G-Biosciences MO, USA) to normalize lipid data. During this procedure, 200 μL of the working solution in 96-well plates was mixed with an aliquot of 25 μL from each replicate’s cell lysate, and the absorbance was measured at 590 nm. Bovine serum albumin (BSA) (G-Biosciences, MO, USA) was diluted serially for standard calibration curve. The slope of the standard calibration curve was then used to compute the protein concentration (mg/mL).

### Statistics and data analysis

Lipid quantification was performed using SCIEX MultiQuant™ with internal standard normalization. Data were adjusted per mg of protein, analyzed from three biological replicates, and assessed for statistical significance (p < 0.05) using Student’s t-test. Graphs were generated with GraphPad Prism 8. PCA plot was created and analyzed using MetaboAnalyst 6.0 software.

## Results

### Experimental evolution of drug-susceptible progenitors to FLC resistance

We previously demonstrated that drug-susceptible and drug-resistant isolates are susceptible to the SL biosynthesis inhibitor Myriocin, compared to *C. albicans* and *C. glabrata*, underscoring the importance of SLs (Kumar *et al*. 2021). In an earlier study, we analysed SL classes and molecular species, revealing a unique molecular profile among FLC-resistant clinical isolates of *C. auris* (Kumar *et al*. 2021). This study investigates how alterations in the SL landscape are associated with the experimental evolution of FLC resistance. For this, two drug-susceptible strains of *C. auris* belonging to clade II were subjected to experimental evolution by exposing them to sub-lethal concentrations of FLC through serial passages over 100 generations (Gerstein and Berman 2020). This led to the development of adapted cells stably displayed a high resistance to the FLC. The adaptation protocol for CBS10913T (C-line) has been previously published by our group Narayanan *et al*. 2022. Similarly, the drug-susceptible isolate NCCPF 470296 (P-line) was also subjected to the same adaptation protocol. After 100 generations, FLC-exposed replicates of both the C-line (C1.1, C1.2 and C1.3) and the P-line (P1.1, P1.2 and P1.3) displayed stable resistance to FLC, with MIC_50_ values ranging from 128 to >256 μg/mL compared to their progenitors (**Supplementary fig. 1a & b**). Single colonies of the replicates showing the highest resistance of >256 MIC_50_ (P1.1 and C1.1) were selected for lipidomic analysis. The drug susceptibility profile of all the adapted replicates are presented in Supplementary Figure 1.

### Drug susceptible C and P-line progenitors displayed distinct SL profiles

The drug-susceptible C and P-line progenitors were also grown separately in a drug-free environment for 100 generations and were used as controls. The SLs from these progenitors were extracted using established protocols and analysed using high-throughput LC-MS/MS techniques, including MRM approaches (Kumar *et al*. 2021). A comparative analysis of SL profiles revealed that both the progenitors of C and P-line (designated as C1 and P1, respectively) contained primary SL intermediates, including LCBs such as DHS, PHS, sphingosine (SPH), and their phosphorylated forms (DHS-1P, PHS-1P, SPH-1P). Other detected intermediates also included phytoceramide (PhytoCer), α-hydroxyphytoceramide (αOHPhytoCer), dihydroceramide (dhCer), ceramide (Cer), α-hydroxyceramide (αOHCer), and α-hydroxylglucosylceramide (αOHGlcCer). The SL intermediates analysis revealed that both the neutral and acidic branches of SL biosynthesis are functional in the C and P-lines. However, significant compositional differences were observed between the progenitors. In C1 control, the most abundant SL class was αOHGlcCer (∼49%), followed by PhytoCer (∼40%), αOHCer (∼5%), dhCer (∼4%) and αOHPCer (∼2%) while LCB (0.54%) and Cer (0.13%) were the least abundant (**Fig. 1a**). In P1 progenitor, PhytoCer (∼53%) was the most abundant SL class followed by αOHPhytoCer (19%). Notably, αOHGlcCer, which was the most abundant class in C1 progenitor, was only ∼12% in P1. This was followed by dhCer (11%) and αOHCer (∼3.5%) whereas similar to C1, LCB (0.49%) and Cer (0.14%) were the least abundant classes in P1 progenitor (**Fig. 1b**). This data indicate massive intra-clade heterogeneity in the SL compositional profiles in both progenitors of the same clade. The clear differences between the two isolates was also confirmed when the data sets were subjected to Principal Component Analysis (**Fig. 1c**)

**Fig. 1:**
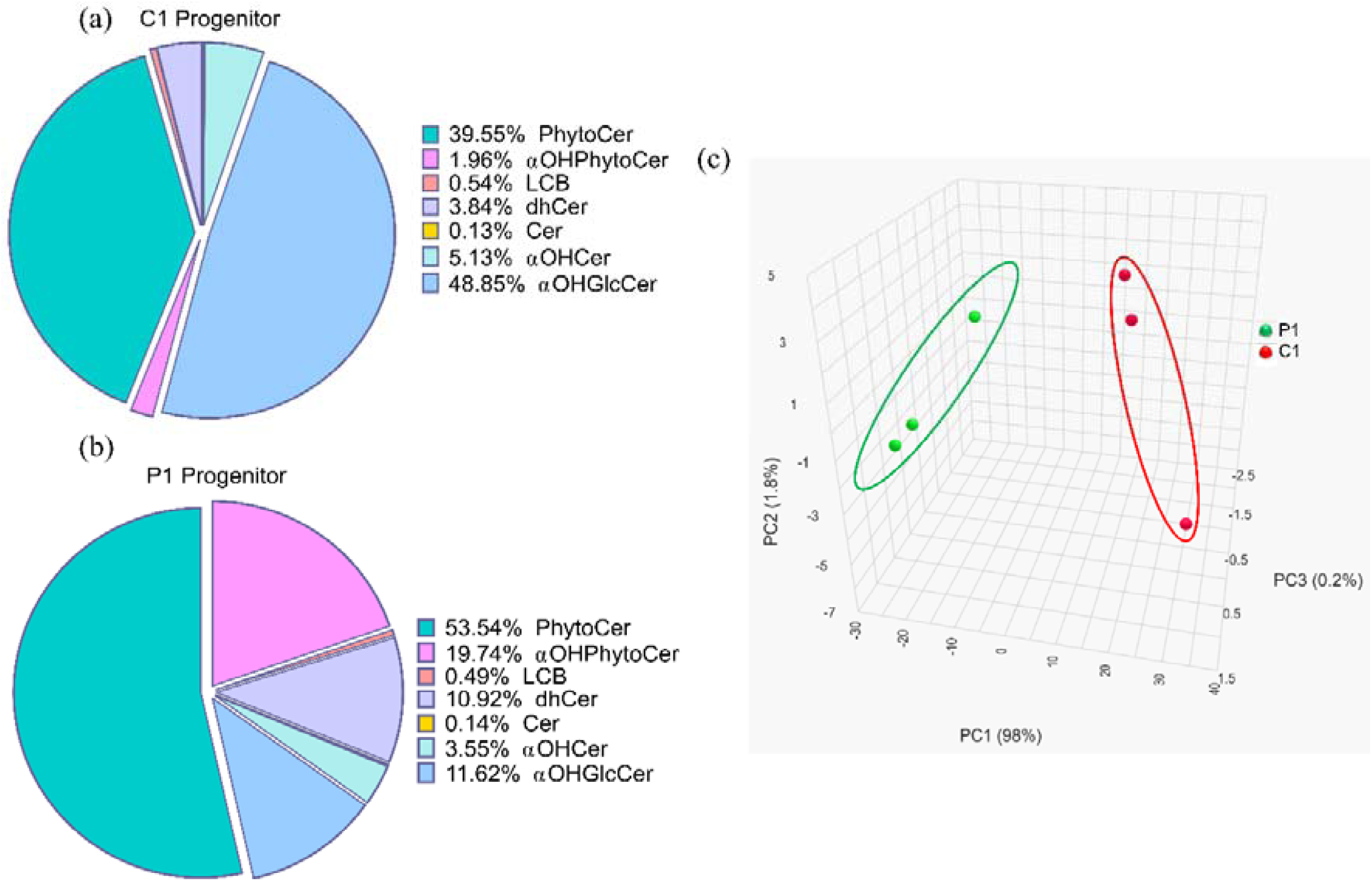
SL compositional differences between the progenitors. **a.** SLs classes distribution in the progenitor C1; **b.** SLs class distribution in the progenitor P1; **c.** Principal component analysis differentiates progenitors on the basis of variations in SL classes between the progenitors.

### Both C and P-line progenitors also exhibit diversity at the SL molecular species level

The molecular species within each SL class, comprising fatty acid chains ranging from 6C to 30C linked to a sphingoid base, vary in carbon chain length, sphingoid backbone structure, and degree of saturation (Kumar *et al*. 2021). Using targeted lipidomics, we focused on key SL molecular species and investigated whether the SL compositional differences between C1 and P1 progenitors were also reflected at the molecular species level. Our analysis revealed that the heterogeneity observed at the SL class levels between C1 and P1 progenitors also extended to the SL molecular species level. For instance, In the C1 progenitor, αOHGlcCer was the most abundant SL class, comprising multiple molecular species. Notably, _GlcCer (d19:2/18:0(OH)) accounted for approximately 49% of the total SL content, making it the single largest molecular species among all the monitored SL species. Other αOHGlcCer species containing 24C (d19:2/24:0(OH)) and 26C (d19:2/26:0(OH)) were present in minor quantities. The neutral SL pathway intermediates, dhCer and Cer, exhibited a molecular species distribution dominated by 24:0, followed by 18:0, 26:0, and 28:0 in both classes. The αOHCer class comprised a diverse range of molecular species with different sphingoid backbones, the most abundant being d18:2 (3.4%), d18:1 (1.4%), and d19:2 (0.05%). Among the LCBs, PHS was the major sphingoid base, constituting 0.49% of the total SLs, followed by DHS and DHS-1-P. Other LCBs, including SPH and SPH-1-P, were detected at very low levels (**Supplementary Fig. 2a**).

In the P1 progenitor, the predominant species in PhytoCer was C24:0, accounting for 37% of the total SLs, followed by 26:0 (12.6%) and a nearly equal amount of C18:0 and C28:0 **(Fig 3b)**. A similar pattern was observed in the αOHPhytoCer class, where the most abundant species was 24:0(OH) (18.1%) and slight contributions from 26:0(OH) (1.5%), 18:0(OH), and 28:0(OH). Notably, in both PhytoCer and αOHPhytoCer classes, the predominant species was the 24:0 in the P1 progenitor. The αOHGlcCer class, representing the third most abundant SL class (12% of total SLs), was primarily composed of two major molecular species: GlcCer (d19:2/18:0(OH) (10.2%) and GlcCer (d18:2/18:0(OH) (1.07%). Similarly, the dhCer class, accounting for ∼11% of total SLs, was predominantly composed of 24:0 dhCer (5%), followed by 18:0 dhCer (4.5%), and minor contributions from 26:0 dhCer, and 28:0 dhCer. The αOHCer class exhibited a distinct molecular profile, with 24:0(OH) (0.66%) as the most abundant species, followed by 26:0(OH) (0.37%), 28:0(OH) (0.30%), and 18:0(OH) (0.23%). Meanwhile, among the LCBs, PHS was the major base (0.39%), and the rest of the molecular species within the LCB class cumulatively contributed 0.48% to total SL content. Lastly, within the Cer class, molecular species were distributed as follows: 24:0 (0.06%), 18:0 (0.026%), 26:0 (0.025%), and 28:0 (0.017%) (**Supplementary Fig. 2b**). These findings highlighted intra-clade heterogeneity extended up to the molecular species level in the C1 and P1 progenitors (**Supplementary Fig. 2a & b**).

### Experimentally evolved resistant adaptors show distinct SL profil*e*

As noted, the drug-susceptible C1 and P1 progenitors displayed distinct SL profiles at the class and molecular species levels. To evaluate how progenitors’ unique SL compositional landscapes influence the evolution of FLC resistance, both progenitors were subjected to experimental evolution. C1.1 adaptor exhibited PhytoCer as the most abundant SL class, accounting for 50% of the total SLs, followed by αOHGlcCer (35.5%), dhCer (6%), αOHCer (∼5%), and αOHPhytoCer (∼3%). The least abundant SL classes in the C1.1 adaptor were LCB (0.42%) and Cer (0.14%) (**Fig. 2a**). Interestingly, αOHGlcCer, a key intermediate in the neutral branch of the SL pathway, decreased by approximately ∼13% of the total SLs in the C1.1 adaptor. This significant reduction was accompanied by a parallel increase in PhytoCer, an intermediate of an acidic branch of the SL pathway implying activation of acidic branch of SL pathway in the C1.1 adaptor. While, dhCer and αOHPhytoCer levels increased modestly along with no significant change in the levels of αOHCer and Cer, the significant reduction in LCB levels was consistent across C1.1 adaptors. These findings highlight distinct alterations in the neutral branch of the SL composition of the C1.1 adaptor, driving a shift from the neutral to the acidic branch of the SL pathway, contributing to FLC resistance **(Fig 2a)**.

**Fig. 2:**
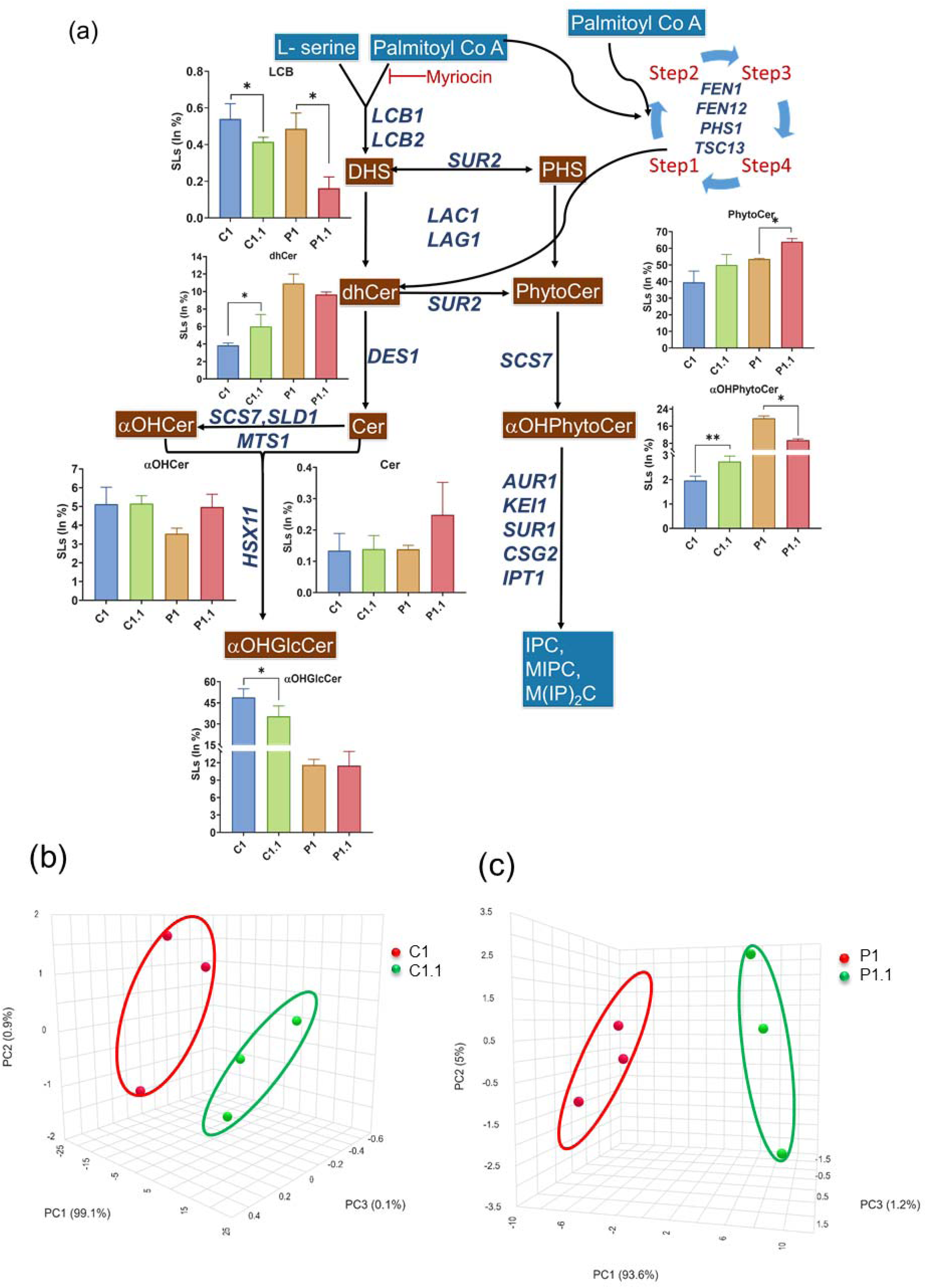
SL profiles of the FLC evolved resistant isolates compared to their progenitor. **a.** Hypothetical SL biosynthesis pathway of *C. auris* and levels of different SL intermediates in each evolved isolates P1.1 and C1.1 as detected by ESI-LCMS/MS. The pathway and genes shown are based on the homologous genes present in *S. cerevisiae*, *C. albicans*. Data on the Y-axis represents % of total SL per mg protein. **b.** 3D PCA plot showing separation of evolved replicates C1.1 from their progenitor C1 based on the compositional differences of SL classes. **c.** 3D PCA plot showing separation of evolved replicates P1.1 from their progenitor P1.

In the P-line, its evolved FLC resistant replicate P1.1 continued to show the dominance of PhytoCer, accounting for > 64% of the total SLs, followed by αOHGlcCer (11%), dhCer (∼10%), αOHPhytoCer (∼9.5%), and αOHCer (∼5%). On the other hand, Cer (0.25%) and LCB (0.16%) were the least abundant SL classes in the P1.1 adaptor. In contrast to the C-line progenitor where αOHGlcCer was the most predominant intermediate, the P-line progenitor had a low level of it, which remained unchanged between P1 and P1.1 cells. (**Fig. 2a**). These findings indicate that αOHGlcCer, the dominant intermediate of a neutral branch of the SL biosynthetic pathway underwent minimal alterations in the P-line isolate during adaptation to FLC resistance. Notably, the intermediates of an acidic branch of the SL pathway exhibited more pronounced changes in P1.1 adaptors. For instance, PhytoCer levels significantly increased in the P1.1 adaptor, while αOHPhytoCer levels were reduced by half of the level of its P1 progenitor (**Fig. 2a**,). This suggests that the acidic branch underwent greater compositional changes in the P-line during FLC adaptation. Interestingly, the modulation of an acidic branch intermediates, PhytoCer and αOHPhytoCer, observed in both C1.1 and P1.1 adaptors was also evident in azole-resistant clinical isolates of *C. auris* (Kumar *et al*. 2021; Ali *et al*. 2024b).

### Principal component analysis (PCA) confirms the statistically significant distinction between P and C-line progenitors and adaptors

PCA was performed using the percentage (% of total SLs) of corresponding datasets, to evaluate statistically significant variations between progenitors C1 and P1 and FLC-evolved isolates of the C1.1 and P1.1. This analysis focused on three major PCA components (PC1, PC2, and PC3) derived from SL class data to capture differences between progenitors. PC1 captures the 98%, PC2 0.2% and PC3 1.8% of the PCA variance. PC1 dominance explains segregation between C1 and P1 based on the SL intermediate PhytoCer, αOHGlcCer and αOHPhytoCer composition **(Fig 1c)**. This variation in the PCA plot were also observed between both lines’ progenitor and FLC evolved adaptors.

For the C-line, the first three components PC1, PC2 and PC3 explained 99%, 0.9%, and 0.1% of the variance, respectively (**Fig. 2b**). The dominance of PC1 indicates a substantial segregation between the C1 progenitor and C1.1 adaptor based on SL class composition. This high proportion underscores the observed variation almost entirely attributable to FLC-induced changes. The increased PhytoCer and decreased GlcCer emerged as the most variable SL classes in the C1.1 adaptor compared to its progenitor. Other SL classes showed minimal variation in the PCA plot associated with evolved FLC resistance. The data points out that the neutral branch of SL metabolism is primarily impacted during the adaptation of FLC resistance in the C-line.

For the P line, PC1, PC2, and PC3 accounted for 93.6%, 5.2%, and 1.2% of the variance, respectively (**Fig. 2c**). These components reflect a broader spread of variation in the P-line compared to the C-line, emphasizing differences in SL class distributions between P1.1 adaptor and P1 progenitor. The key contributors to variation were increased PhytoCer and decreased aOHPhytoCer levels in the P1.1 adaptor. Other SL classes demonstrated minimal changes between the P1 progenitor and P1.1 adaptor, suggesting that FLC-induced resistance in the P line primarily affects the acidic branch of SL metabolism.

Overall, PCA effectively illustrates the statistically significant impact of FLC induced resistance on SL fingerprints in both line isolates. These findings suggest strain-specific SL remodelling during the evolution of FLC resistance and also provide insights into intra-clade SL class-specific contributions to the adaptive response of *C. auris* under FLC pressure.

### FLC-evolved resistant lines show significant alterations at the molecular species level

We analyzed the molecular species composition of the drug-evolved resistant isolates P1.1 and C1.1 and compared them with their respective progenitors P1 and C1. Alongside overall compositional differences, distinct changes in the distribution of molecular species within the same SL class were also observed. In the C1.1 adaptor, the predominant molecular species within the PhytoCer class were C24:0 and C26:0, wherein only C24:0 levels showed a significant increase compared to the C1 progenitor (**Fig. 3a**). A similar trend was observed in αOHPhytoCer class, wherein C24:0(OH) species was the only molecular species that exhibited a significant increase while other species remained unchanged (**Fig. 3a & b**). These compositional alterations indicate that 24C fatty acyl chains of PhytoCer and αOHPhytoCer were preferentially enriched in the C1.1 adaptor during the drug evolution process.

**Fig. 3:**
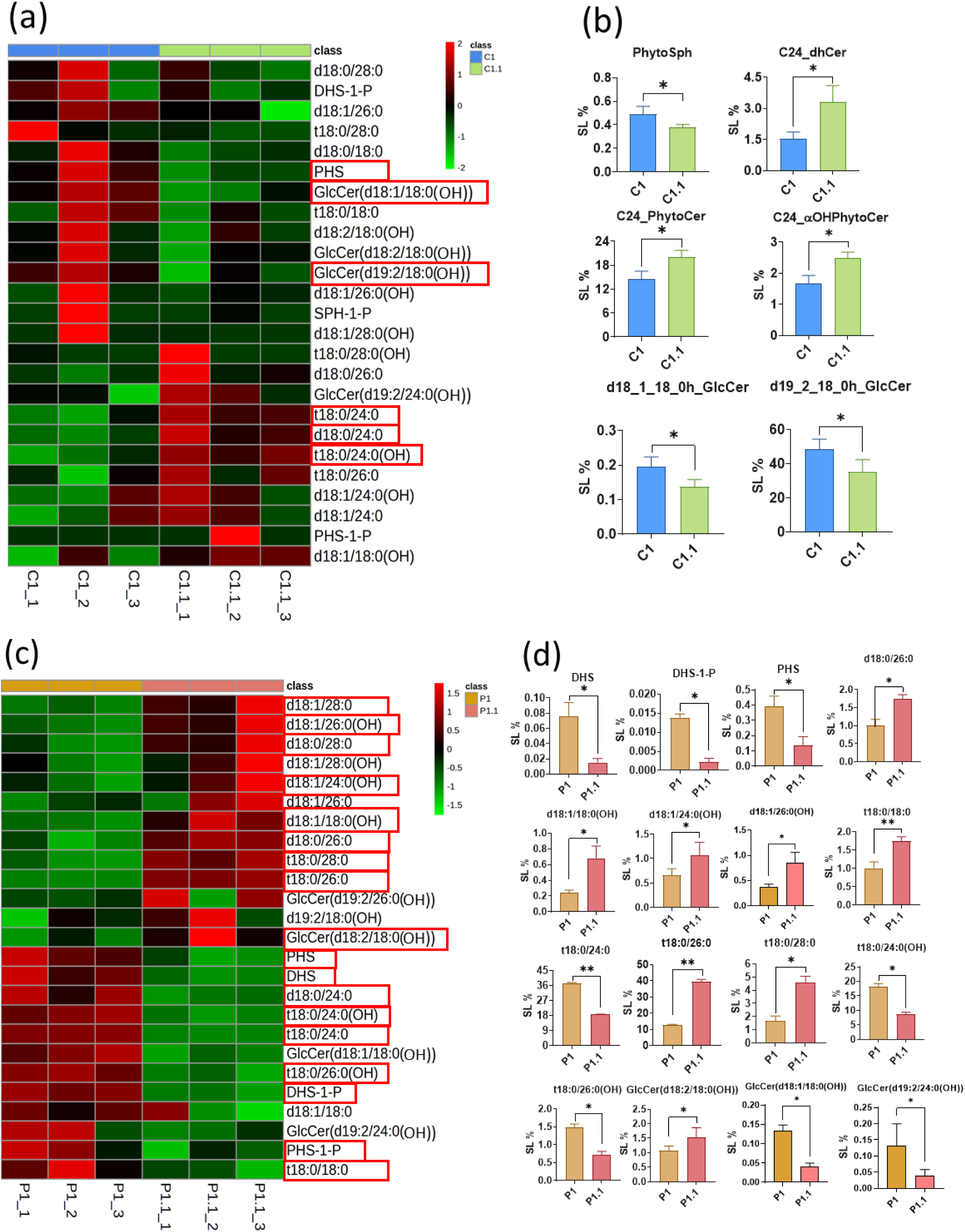
Major SL molecular species of FLC evolved resistant isolates of both lines compared to their progenitor. **a**. Heat map depicting the variation of SL molecular species of C1.1 evolved replicates and progenitor C1. Lipid contents were normalized and plotted by MetaboAnalyst 6 **b.** Significantly altered molecular species between evolved C1.1 and progenitor C1 were quantified and plotted using Graph Pad Prism. **c.** Heat map depicting the variation of SL molecular species of P1.1 evolved replicates and progenitor P1. Lipid contents were normalized and plotted by MetaboAnalyst 6. **d.** Significantly altered molecular species between evolved P1.1 and progenitor P1 were quantified and plotted using Graph Pad Prism. Significance was checked using Student’s t-test(* *p*<0.05, ** *p*<0.01)

The second most abundant SL class in the C1.1 adaptor was αOHGlcCer, which accounted 35.55% of the total SL composition, significantly lower than its progenitor C1. Among the two major molecular species of αOHGlcCer, the predominant species, GlcCer d19:2/18:0(OH), accounted for almost all of the αOHGlcCer content i.e.; 35% of the total SL content in the C1.1 adaptor and was significantly reduced from 48.3% in the progenitor C1 (**Fig. 3a & b**). The second molecular species, GlcCer d18:1/18:0(OH), was low in abundance while other molecular species within this class remained unchanged, indicating specific and targeted alteration of d19:2/18:0(OH) composition rather than a broad restructuring of the entire class. These findings suggest that the reduction in αOHGlcCer levels, particularly the depletion of the dominant GlcCerd19:2/18:0(OH) species, maybe a key aspect of the adaptive response in C1.1. In addition, the major LCB, PHS-the first intermediate in the t18:0 branch was reduced in C1.1 compared to C1, suggesting that PHS was actively fluxed to synthesis of PhytoCer as evident from the increased content of the latter in the adaptor (**Fig. 3a & b**). Other species of dhCer and LCB showed no significant changes in the C-line adaptor.

In the P1.1 adaptor, a significant increase in the PhytoCer class was also observed. In contrast to C1.1, C26:0 emerged as the dominant species that exhibited a remarkable increase (12.6% in P1 to 39.3% in P1.1), replacing C24:0, which was the most abundant in P1 (**Fig. 3c & d**). A similar trend was noted for C28:0, which increased from 1.7% in P1 to 4.5% in P1.1.

In contrast, C24:0 and C18:0 PhytoCer levels declined from 37.5% and 1.6% in P1 to 18.6% and 1.2% respectively in P1.1. These changes highlight a distinct shift in PhytoCer composition, favouring PhytoCer with much longer acyl chains (26:0 and 28:0) in the P1.1 adaptor over 18:0 and 24:0 (**Fig. 3c & d**). A peculiar observation was seen in the αOHPhytoCer which is the immediate downstream product of PhytoCer. More than 10-fold reduction was observed in C24:0(OH) and C26:0(OH) species of αOHPhytoCer in the adaptor while C18:0(OH) and C28:0(OH) levels remained unchanged (**Fig. 3c & d**). These compositional changes suggest heavy suppression of PhytoCer to αOHPhytoCer conversion as an adaptive response influencing membrane composition at the molecular level in P1.1. Interestingly, overall αOHGlcCer remained unchanged at the class level; only a single species d18:2/18:0(OH) was slightly increased. The dhCer class showed no major alterations, though C24:0 dhCer levels decreased from 5% to 3%, while C26:0 dhCer rose from 1% to 1.7% in P1.1. Other dhCer species exhibited no significant variations. Among LCB molecular species, PHS, the dominant sphingoid base, showed a marked reduction, declining from 0.39% in P1 to 0.13% in P1.1 (**Fig. 3c & d**), an observation also seen in C1.1. Notably, molecular species of the αOHCer class ranging from C18 to C28 showed an increasing trend, but only C18:0(OH) significantly increased in P1.1 (**Fig. 3c & d**). The Cer class exhibited no significant changes at the molecular species level in the P1.1 adaptor.

### Alteration of Sterol Composition in Fluconazole-Resistant Adaptors

Ergosterol is an essential component of fungal cell membranes, functioning analogously to cholesterol in mammalian cells. Azoles inhibit ergosterol biosynthesis by targeting lanosterol 14α-demethylase (*ERG11/CYP51*) (Moye-Rowley 2020),compromising fungal membrane integrity and function. To further investigate the impact of FLC resistance, we analysed the sterol composition of FLC-adapted isolates, which had already demonstrated reprogramming of SL composition during experimental evolution (discussed above). We examined the sterol composition of C1 and P1 progenitors and compared them with their respective adapted isolates (C1.1 and P1.1) using Gas Chromatography-Mass Spectrometry (GC-MS), as described by Carolus *et al*(Carolus *et al*. 2024). Our GC-MS analysis identified key intermediates in the sterol biosynthesis pathway, accounting for most of the total sterol content. Consistent with previous reports, ergosterol was the predominant sterol in *C. auris*, comprising more than 80% of the total cellular sterols. Other sterol intermediates, including lanosterol, eburicol, fecosterol, episterol, zymosterol, ergosta-5,7-dienol, cholesta-7,24-dienol, cholesta-5,7,24-trienol, and cholesta-5,7,22,24-tetraenol, were present in lower amounts **(Fig. 4a)**.

**Fig. 4:**
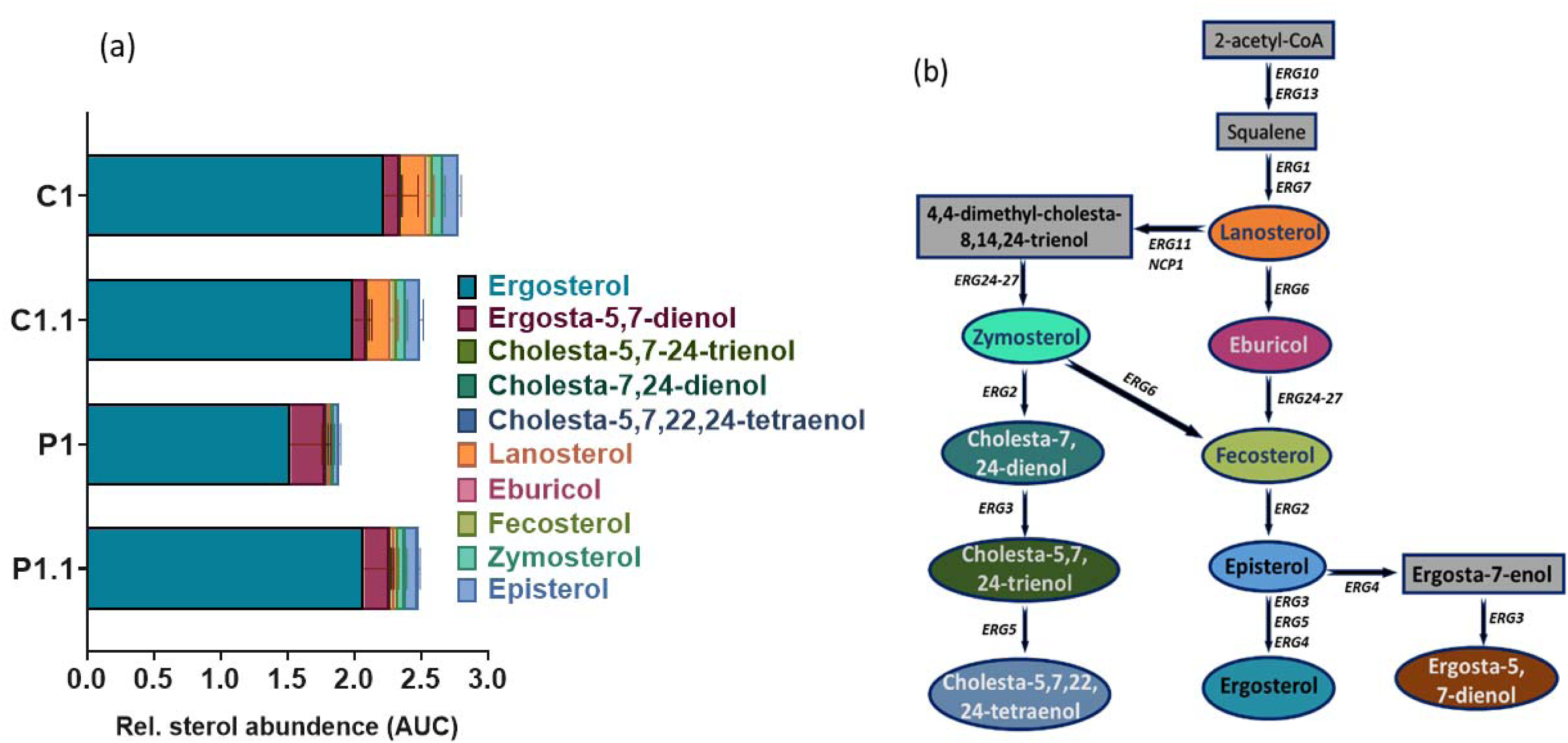
Sterol compositional changes in C- and P-lines. **a.** The bar plot presents GC-MS-based sterol abundance, measured as the area under the curve (AUC) relative to the internal standard control under drug-free conditions in both the adaptor and progenitor. **b.** The predicted ergosterol biosynthesis pathway is shown, with colored intermediates corresponding to the intermediate levels in (a). All experiments were conducted in biological triplicates, and the means and standard deviations were calculated.

Notably, the C-line progenitor C1 exhibited a significantly higher total sterol content than its adapted strain, C1.1. As observed, ergosterol was the predominant sterol (>80%) in the C-line progenitor, followed by lanosterol, ergosta-5,7-dienol, and episterol, while other sterol intermediates were present in lower amounts. Notably, after 100 generations of FLC exposure, total sterol content was reduced in the C1.1 adaptor, specifically driven by a decline in ergosterol levels, without affecting the concentrations of other sterol intermediates. This decline in ergosterol aligns with the known azole mechanism of action, which inhibits the ergosterol biosynthesis pathway by targeting the lanosterol 14α-demethylase enzyme (*ERG11*). These findings suggest that prolonged FLC exposure led to a selective reduction in ergosterol levels in the C1.1 adaptor.

In contrast, the P1.1 adaptor exhibited a more than 20% increase in total sterol content compared to its progenitor, P1. However, ergosterol remained the predominant sterol in both P1 and P1.1 cells, followed by ergosta-5,7-dienol, while other sterol intermediates were present in relatively lower amounts. Upon FLC exposure, the increase in total sterol content in the P1.1 adaptor was primarily driven by elevated ergosterol levels, with no significant changes observed in other sterol intermediates, making ergosterol the most abundant sterol (**Supplementary file 1**). This finding aligns with a previous study which reported a similar increase in ergosterol content in *C. auris* cells following azole treatment (Rybak *et al*. 2020). In contrast, ergosta-5,7-dienol levels slightly reduced in the P1.1-adaptor **(Figure 4a & b)**. These results indicate that ergosterol was the only sterol that underwent significant compositional and quantitative changes in response to FLC exposure. In contrast, the levels of other sterols remained unchanged in the P1.1 FLC-adaptor. Although, sterol compositional changes in experimentally evolved FLC-resistant strains are recorded in response to experimental evolution between the progenitors under antifungal stress. The results also indicate that FLC resistance is not solely dependent on sterols but may also involve changes in other membrane lipids contributing to drug resistance.

## Discussion

FLC resistance is influenced by multiple factors, including the overexpression of drug efflux pumps, mutations in *ERG* genes, stress response pathways, biofilm formation, and cell wall integrity, underscoring its multifaceted nature. While these canonical azole resistance mechanisms are well-documented in *C. auris*, they fail to fully explain the unprecedented levels and frequency of azole resistance exhibited by this pathogen. This gap in understanding has driven research toward identifying novel mechanisms and targets that may play a crucial role in *C. auris* drug resistance. Recent studies, including those from our group, suggest that SLs composition can modulate membrane fluidity and drug interactions independent of ergosterol content, thereby influencing azole resistance. The present study strengthens the case for SLs as key determinants of multidrug resistance (MDR) in *C. auris*. This conclusion is supported by an in-depth SL analysis of two independently evolved *C. auris* clade II isolates that developed stable FLC resistance.

The observed alterations in SL profiles suggest strain-specific remodelling strategies that favouring either the neutral or acidic branches of the SL biosynthesis pathway. For example, the P-line (P1) progenitor strain appears to favour the acidic branch, as indicated by elevated levels of PhytoCer, and αOHPhytoCer. In contrast, the C-line progenitor (C1) predominantly relies on the neutral branch, characterized by increased levels of long-chain bases (LCBs), αOHCer, and αOHGlcCer (**Fig. 4**). The intra-clade heterogeneity in SL profiles among progenitor strains highlights intrinsic metabolic differences that may predispose isolates to distinct resistance trajectories. While P1 exhibited a dominance of 24:0 PhytoCer, C1 favoured αOHGlcCer, pointing to lineage-specific SL metabolic programs.

Certain commonalities emerged despite the SL compositional heterogeneity between the two progenitors and their evolved lines. Notably, an increased abundance of PhytoCer was consistently associated with fluconazole (FLC) resistance in both independently evolved adaptors. Principal component analysis (PCA) further validated the distinct SL landscape in FLC-evolved adaptors, with PC1 capturing high variance in both C- and P-line isolates—underscoring SL remodelling as a key factor in resistance acquisition. The elevated PhytoCer levels and reduced αOHPhytoCer in P1.1 emphasize the selective pressure on the acidic SL pathway during FLC adaptation (**Fig. 4**). Similarly, C1.1 displayed a parallel pattern but with a more pronounced reduction in αOHGlcCer, suggesting differential metabolic rewiring between isolates. The increased PhytoCer levels alongside the decrease in αOHGlcCer at both SL class and molecular species levels point to the modulation of acidic SLs, a feature also observed in FLC-resistant clinical isolates of *C. auris* from clades I, III, and VI (Kumar *et al*. 2021; Ali *et al*. 2024b). This convergence highlights the relevance of acidic SLs in resistance phenotypes. Acidic SLs play critical roles in yeast physiology, including membrane organization, signal transduction, and stress responses. In the context of antifungal resistance, these lipids contribute to membrane rigidity, drug efflux regulation, and stress adaptation (Prasad et al. 2005; Shahi et al. 2022). SL composition changes can affect multidrug transporter function and influence yeast survival under antifungal pressure. Additionally, acidic SLs participate in pathways such as the Pkh1/2-Ypk1 signalling cascade, which regulates membrane homeostasis and the antifungal response (Víglaš and Olejníková 2021). Our findings further support that the modulation of acidic SLs which are absent in human hosts are unique to *Candida* species which represents a key adaptive strategy in experimentally evolved isolates, underscoring their significance in antifungal resistance.

While sterol homeostasis is a key factor in azole resistance, substantial evidence indicates that resistance can also arise independently of sterol modulation (Lee, Robbins and Cowen 2023). In our study, FLC-evolved strains and their progenitors displayed largely similar sterol profiles, whereas the adapted strains exhibited significant reprogramming of their SL composition (**Fig 5**). These findings underscore the central role of SLs in the evolution of FLC resistance in *C. auris*, consistent with observations in clinical FLC-resistant isolates. Although our data strongly support the involvement of SLs in FLC resistance, it is essential to recognize the study’s limitations, as experiments were conducted using only two independent drug-susceptible progenitors and a single antifungal, FLC, for experimental evolution. Therefore, broader conclusions require further validation across a more diverse set of isolates and triazole antifungals.

**Fig. 5:**
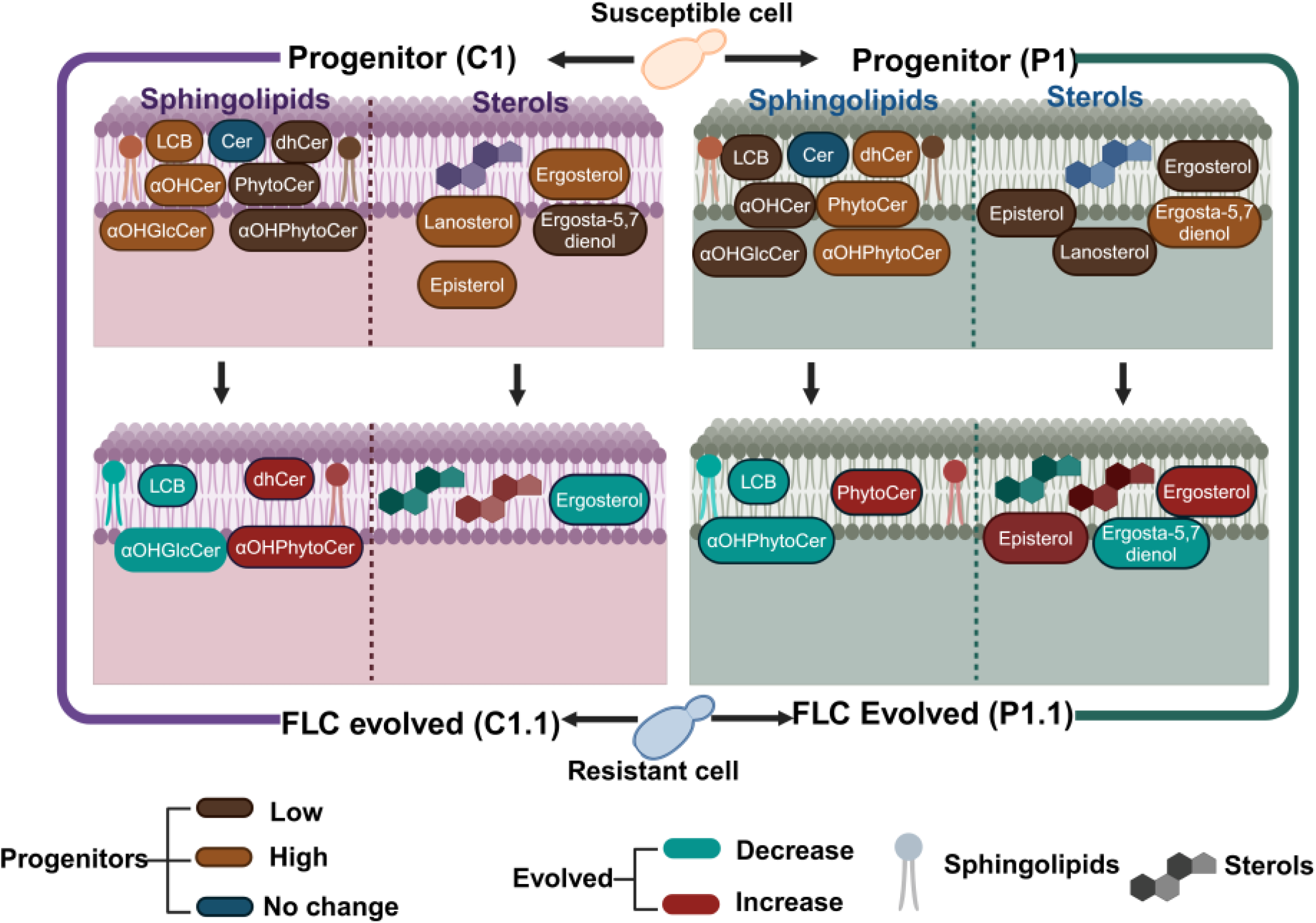
Schematic depiction of SLs and sterol composition changes in two clade II progenitors of C. auris and their evolved FLC-resistant variants.

## Acknowledgments

This study was funded by the Indian Council of Medical Research (AMR/149/2018-ECD-II), the Government of India, to R.P., K.S., A.C., and S.M.R. R.P. acknowledges the Department of Biotechnology support under Boost to University Interdisciplinary Life Science Departments for Education and Research Programme (DBT-Builder), and DBT-PG TEACHING support to Amity Institute of Biotechnology, Amity University Haryana. P.K. acknowledges the Indian Council of Medical Research for the award of a Senior Research fellowship (2020-7196/CMB-BMS). B.A. acknowledges the Council of Scientific and Industrial Research fellowship (09/263(1223)/2019-EMR-1). A.C. acknowledges the Indian Council of Medical Research fellowship (Myco/Fell/7/2022-ECD-II). M.K. acknowledges the award of ICMR-Research Associateship Fellowship (Myco/Fell/3/2022-ECD-II). P.K., B.A., and A.C. acknowledge the support of the Amity Central Instrument Research facility (CIRF) and Amity Lipidomics Research Facility (ALRF) for carrying out this work.

---

**Supplementary figure 1:**
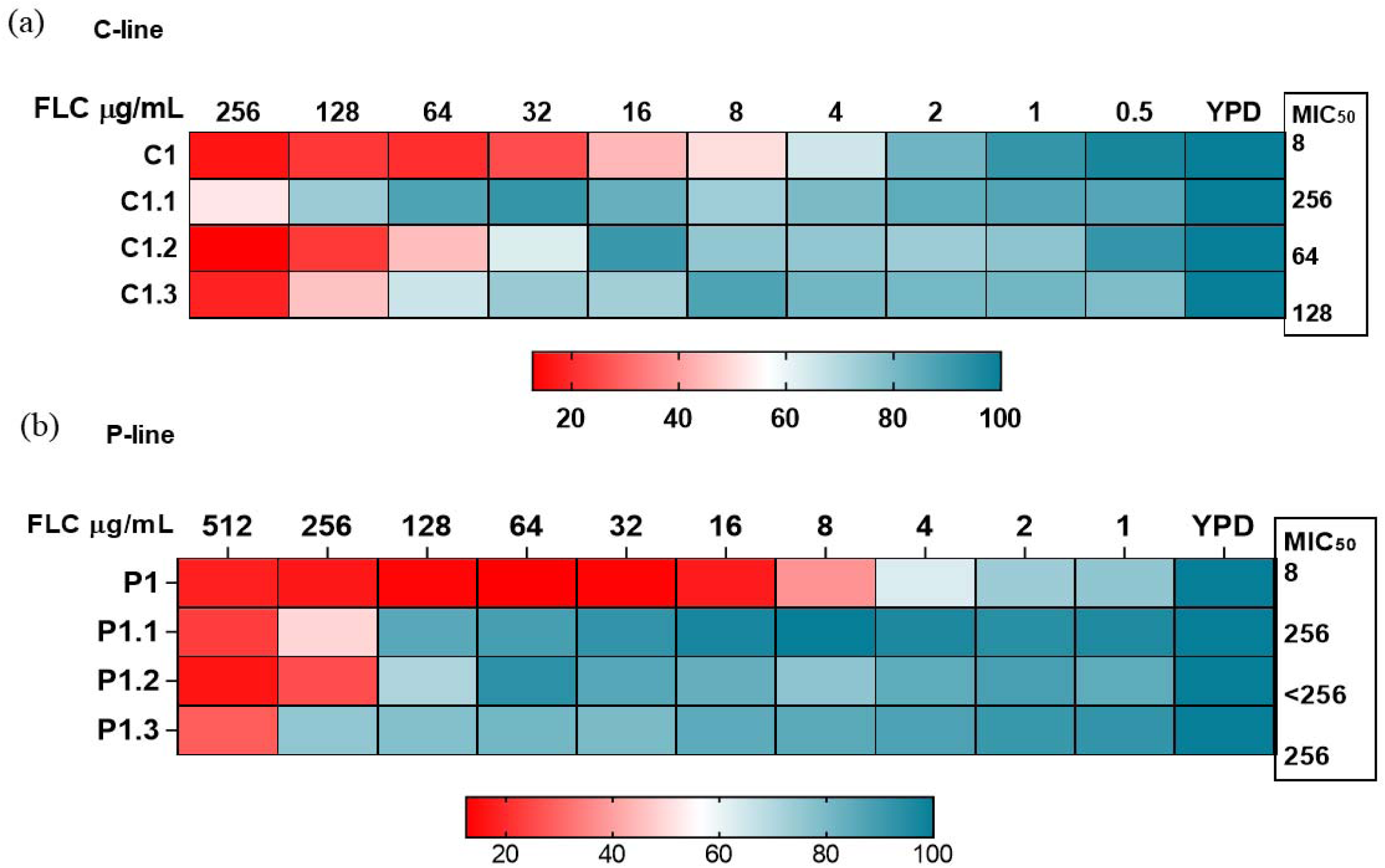
YPD media was used to culture *C. auris* cells at 30°C for overnight, and MIC tests were carried out in accordance with Clinical and Laboratory Standards Institute (CLSI) recommendations. **a.** Microdilution assay was conducted for the successful evolved FLC resistance of C-line progenitor and evolved replicates *C. auris*. **b.** Microdilution assay was conducted for the successful evolved FLC resistance of P-line progenitor and evolved replicates of *C. auris*.

**Supplementary figure 2:**
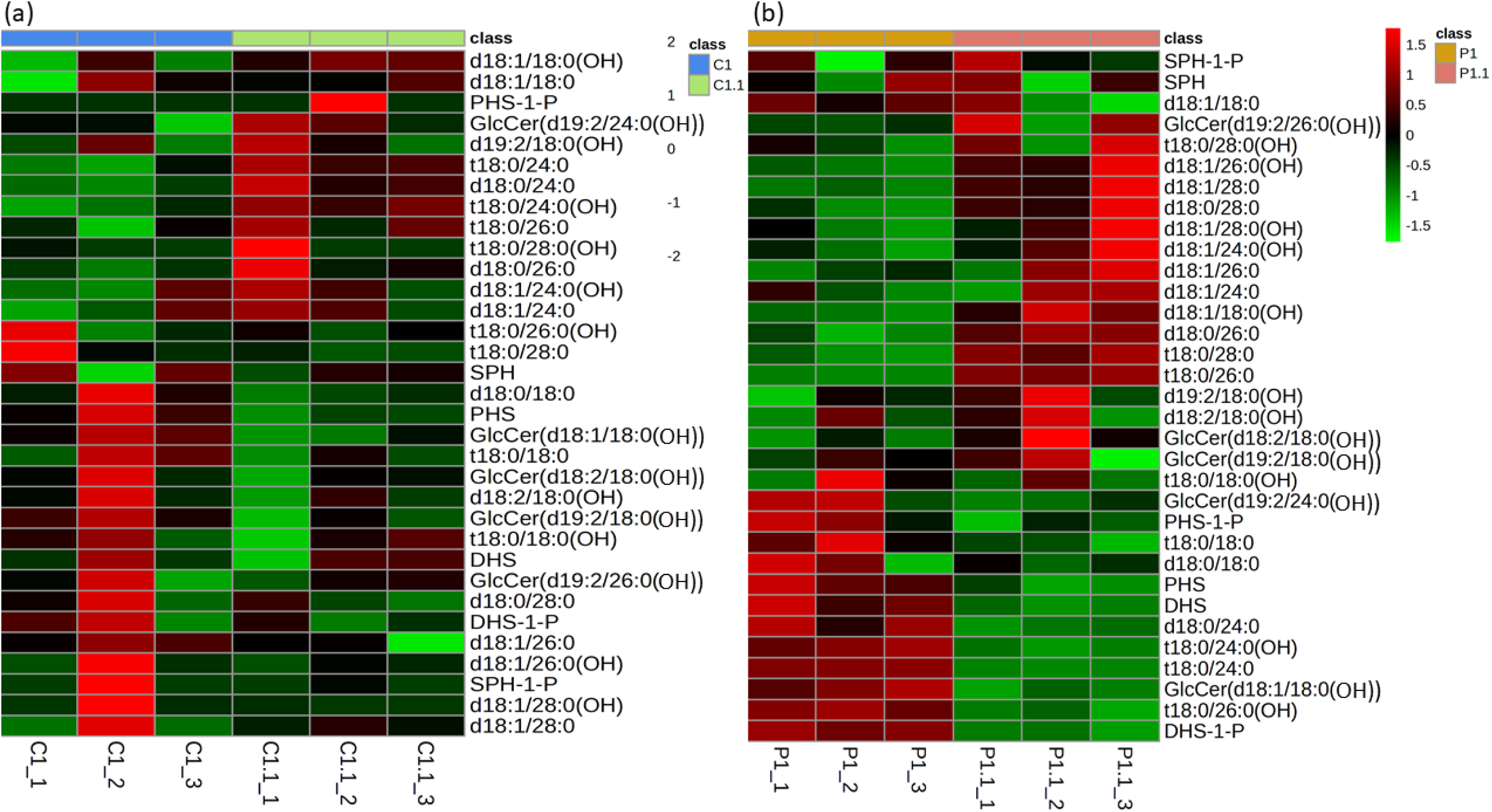
SL molecular species of FLC evolved resistant isolates of both line compared to their progenitor. **a**. Heat map depicting variation of total SL molecular species of P1.1 evolved replicates and progenitor P1. Lipid contents were normalized and plotted by MetaboAnalyst 6 **b.** Heat map depicting variation of total SL molecular species of C1.1 evolved replicates and progenitor P1. Lipid contents were normalized and plotted by MetaboAnalyst 6.

